# A Rapid Multiplex LAMP Assay for Point-of-Care Detection of CT, NG, TV, and Fluoroquinolone Resistance in NG

**DOI:** 10.1101/2025.07.24.666671

**Authors:** Shoulian Dong, Manuel X. Duval, Tri D. Do, Jeff Ho, Omar Abdallah, Michael R. White, Jacob C. Johnson, Christine Meda, Nancy Schoenbrunner

## Abstract

Rapid point-of-care (POC) diagnostics are essential tools for improving timely treatment and reducing the transmission of sexually transmitted infections (STIs). The STI NG Plus Assay is a rapid multiplex LAMP (loop-mediated isothermal amplification) NAAT (nucleic acid amplification test) capable of simultaneously detecting *Chlamydia trachomatis* (CT), *Neisseria gonorrhoeae* (NG), *Trichomonas vaginalis* (TV), and fluoroquinolone resistance-associated mutations in NG (*gyrA* S91F). In this study, we assessed the STI NG Plus assay primer design and analytical sensitivity. Using a bioinformatically optimized primer design pipeline and empirical screening, the assay demonstrated high inclusivity and specificity, with no cross-reactivity to 48 urogenital organisms or the human genome. Analytical sensitivity testing showed reliable detection of all targets in both lysis buffer and clinical matrix. Limits of detection were lower than those of an existing FDA-cleared test. The assay’s robustness, speed, and sensitivity support its potential for decentralized STI testing with integrated antimicrobial resistance profiling.

## INTRODUCTION

Rapid diagnostic tests at the point-of-care (POC) are transforming infectious disease management by significantly reducing the time to diagnosis and treatment initiation, thus improving patient outcomes. Timely, accurate diagnostics in clinical settings enable immediate treatment decisions, enhance patient compliance, and reduce disease transmission, particularly in sexually transmitted infections (STIs) where follow-up visits may be challenging [1,2]. POC testing can enable a shift from empirical, symptom-based or history-based treatment to timely, result-driven treatment, reducing unnecessary exposure to antibiotics. Adoption of POC diagnostics has the potential to reduce total cost of care, when implemented in an evidence-based manner [3-5]. Studies have demonstrated that POC testing leads to improved patient satisfaction, reduced patient anxiety, and better clinical outcomes due to faster initiation of treatment [6,7].

A critical and growing threat within STI management is antimicrobial-resistant (AMR) *Neisseria gonorrhoeae* (NG). The World Health Organization (WHO) has classified NG as a high-priority pathogen due to escalating resistance to multiple antibiotic classes, including fluoroquinolones and extended-spectrum cephalosporins [8,9]. Molecular assays capable of detecting fluoroquinolone resistance mutations, such as the *gyrA* S91F mutation, provide an opportunity for targeted antibiotic therapy. Such resistance-informed testing promotes antibiotic stewardship by reducing inappropriate antibiotic use and preserving the effectiveness of existing antibiotic therapies [10-12].

Multiplex diagnostic platforms further enhance clinical utility by simultaneously detecting multiple pathogens and resistance markers. A multiplex POC assay capable of identifying *Chlamydia trachomatis* (CT), *Neisseria gonorrhoeae* (NG), *Trichomonas vaginalis* (TV), and fluoroquinolone resistance mutations in NG will offer substantial clinical advantages. Enhanced sensitivity, particularly in assays utilizing loop-mediated isothermal amplification (LAMP), increases diagnostic accuracy, ensuring fewer false negatives and thus enabling more reliable clinical decision-making [13-15]. This improvement directly impacts patient outcomes through timely and appropriate interventions, reduced complications from untreated infections, and improved long-term reproductive health [16,17].

Loop-mediated isothermal amplification (LAMP) technology has emerged as a promising alternative NAAT to traditional quantitative polymerase chain reaction (qPCR) due to several distinct advantages. LAMP is rapid, simple, and cost-effective, requiring minimal laboratory equipment and enabling use in resource-limited settings, making it ideal for point-of-care testing [18-20]. Moreover, LAMP assays often demonstrate higher tolerance to inhibitors commonly found in clinical samples, enhancing their robustness in field applications [21].

However, LAMP has certain limitations compared to qPCR, including potential difficulties in multiplexing due to complex primer design, reduced quantitative capability, and occasional challenges in distinguishing specific amplification products from nonspecific signals [22,23]. These limitations make careful assay design exceedingly important. For molecular diagnostic assays, performance is highly dependent on the specificity and inclusivity of the primers and probes. Inclusivity ensures that all known genetic variants of a target organism are reliably detected, while specificity ensures that the assay does not cross-react with closely related organisms, other microbes present at the sample collection site, or human genomic DNA. These considerations are particularly critical for LAMP, which uses multiple primers per target and is thus more prone to primer-dimer formation and off-target amplification. The STI NG Plus assay was therefore designed using a rigorous *in silico* pipeline to ensure comprehensive genome coverage and minimal cross-reactivity, followed by empirical screening to confirm the absence of primer-primer interference and the robustness of each target’s detection chemistry.

In this study, we present early sensitivity data from an extraction-free multiplex LAMP-based assay targeting CT, NG, TV, and NG *gyrA* mutations associated with fluoroquinolone resistance, in addition to an *in silico* analysis of LAMP primers used. The assay’s chemistry and analytical sensitivity are evaluated, demonstrating its suitability as a reliable diagnostic tool that may be used in diverse clinical settings. While this initial testing was carried out using a research instrument, the assay is designed for use in point-of-care settings on a device that can be operated by untrained individuals with minimal instruction.

## MATERIALS/METHODS

### *In Silico* Assay Design and Bioinformatic Screening

Candidate LAMP primer sets were designed for each target gene using publicly available genomic sequence data. To assess inclusivity, full-length genome sequences for each pathogen (*C. trachomatis, N. gonorrhoeae*, and *T. vaginalis*) were retrieved from the NCBI Nucleotide database via the Entrez API. For each target, local pairwise alignments were performed between candidate primers and all retrieved genome records using the Bioconductor Biostrings package in R. Primer sets were retained only if all oligonucleotides aligned to the complete set of known isolates with minimal mismatches (≤1 mismatch per primer), ensuring detection across all known variants.

Specificity and cross-reactivity were then evaluated against a panel of 48 organisms commonly found in the urogenital tract (**Table 1)**, including bacteria, fungi, viruses, and protozoa. Local alignments were performed to identify potential off-target matches at genomic loci that could support amplification. Primer sets with significant cross-reactivity—defined as alignment of multiple primers within a set to a single non-target genome—were excluded. All primers were also aligned against the human genome (GRCh38.p13) to exclude any potential interaction with host DNA. Primer sets passing both inclusivity and specificity filters were selected for empirical validation. A schematic of the bioinformatic screening pipeline is shown in **(Figure 1)**.

**Table 1.** List of organisms included in cross-reactivity panel. This panel includes organisms known to colonize or infect the urogenital tract. Sequence data were retrieved from the NCBI database and used for local alignment-based cross-reactivity *in silico* screening.

| Organism | Type |
| --- | --- |
| *Fannyhessea vaginae* | Bacterium |
| *Bacteroides fragilis* | Bacterium |
| *Campylobacter ureolyticus* | Bacterium |
| *Bifidobacterium longum* | Bacterium |
| *Candida albicans* | Fungus |
| *Nakaseomyces glabratus* | Fungus |
| *Candida parapsilosis* | Fungus |
| *Chlamydia pneumoniae* | Bacterium |
| *Chlamydia psittaci* | Bacterium |
| *Chlamydia trachomatis* | Bacterium (non-target strain) |
| *Clostridium perfringens* | Bacterium |
| *Corynebacterium genitalium* | Bacterium |
| *Corynebacterium xerosis* | Bacterium |
| *Enterococcus faecalis* | Bacterium |
| *Escherichia coli* | Bacterium |
| *Gardnerella vaginalis* | Bacterium |
| *Haemophilus ducreyi* | Bacterium |
| *Human alphaherpesvirus 1* | Virus |
| *Human alphaherpesvirus 2* | Virus |
| *Human papillomavirus* | Virus |
| *Kingella denitrificans* | Bacterium |
| *Kingella kingae* | Bacterium |
| *Klebsiella pneumoniae* | Bacterium |
| *Lactobacillus acidophilus* | Bacterium |
| *Levilactobacillus brevis* | Bacterium |
| *Lactobacillus jensenii* | Bacterium |
| *Lactobacillus delbrueckii* subsp. *lactis* | Bacterium |
| *Megasphaera lornae* | Bacterium |
| *Mobiluncus curtisii* | Bacterium |
| *Mobiluncus mulieris* | Bacterium |
| *Moraxella lacunata* | Bacterium |
| *Mycoplasma genitalium* | Bacterium |
| *Metamycoplasma hominis* | Bacterium |
| *Bergeriella denitrificans* | Bacterium |
| *Neisseria gonorrhoeae* (non-target strain) | Bacterium |
| *Neisseria meningitidis* | Bacterium |
| *Neisseria subflava* | Bacterium |
| *Peptostreptococcus anaerobius* | Bacterium |
| *Proteus mirabilis* | Bacterium |
| *Pseudomonas aeruginosa* | Bacterium |
| *Staphylococcus aureus* | Bacterium |
| *Staphylococcus epidermidis* | Bacterium |
| *Streptococcus agalactiae* | Bacterium |
| *Trichomonas vaginalis* (non-target strain) | Protozoan |
| *Ureaplasma parvum* | Bacterium |
| *Ureaplasma urealyticum* | Bacterium |
| *Shigella sonnei* | Bacterium |
| *Shigella dysenteriae* | Bacterium |
| *Homo sapiens* (GRCh38) | Host genome |

**Figure 1.**
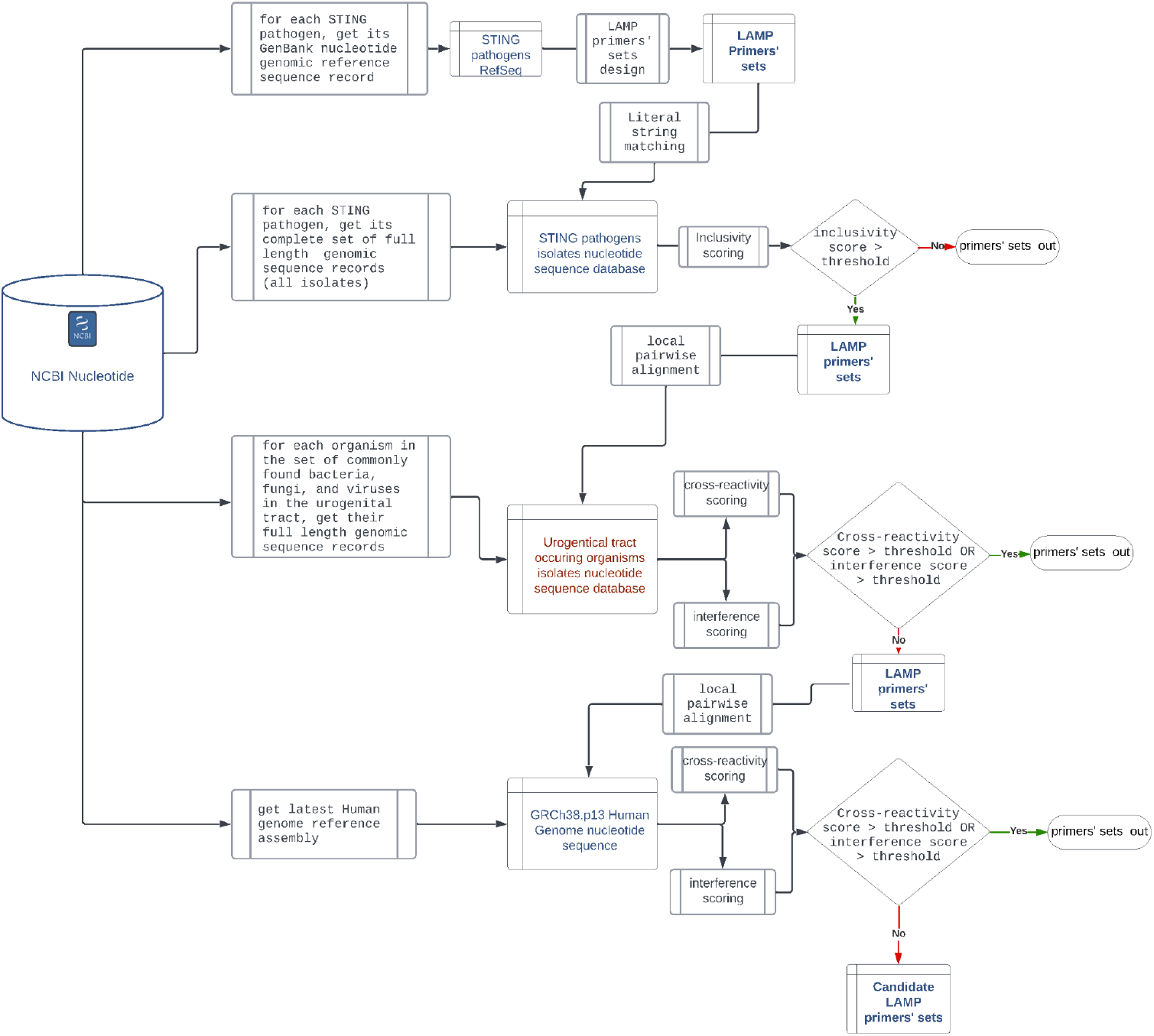
*In silico* primer screening workflow for STI NG Plus assay design. The bioinformatics pipeline used for LAMP primer design and validation involved three sequential screening stages: (1) inclusivity assessment to ensure primer binding across all known genome variants of each target pathogen; (2) cross-reactivity assessment against 48 non-target organisms commonly found in the urogenital tract; and (3) host genome screening to eliminate any potential interaction with human genomic DNA. Only primer sets passing all three filters were advanced to laboratory screening as shown in Figure 2.

### Empirical Screening of Primer Sets

Primer sets that passed *in silico* screening were synthesized and evaluated in the laboratory (**Figure 2**). Initial screening was performed in singleplex LAMP reactions using synthetic double-stranded gene fragments corresponding to each target. Reactions were monitored using an intercalating dye to measure time-to-detection (TTD), with a TTD of ≤15 minutes under screening conditions required to proceed.

**Figure 2.**
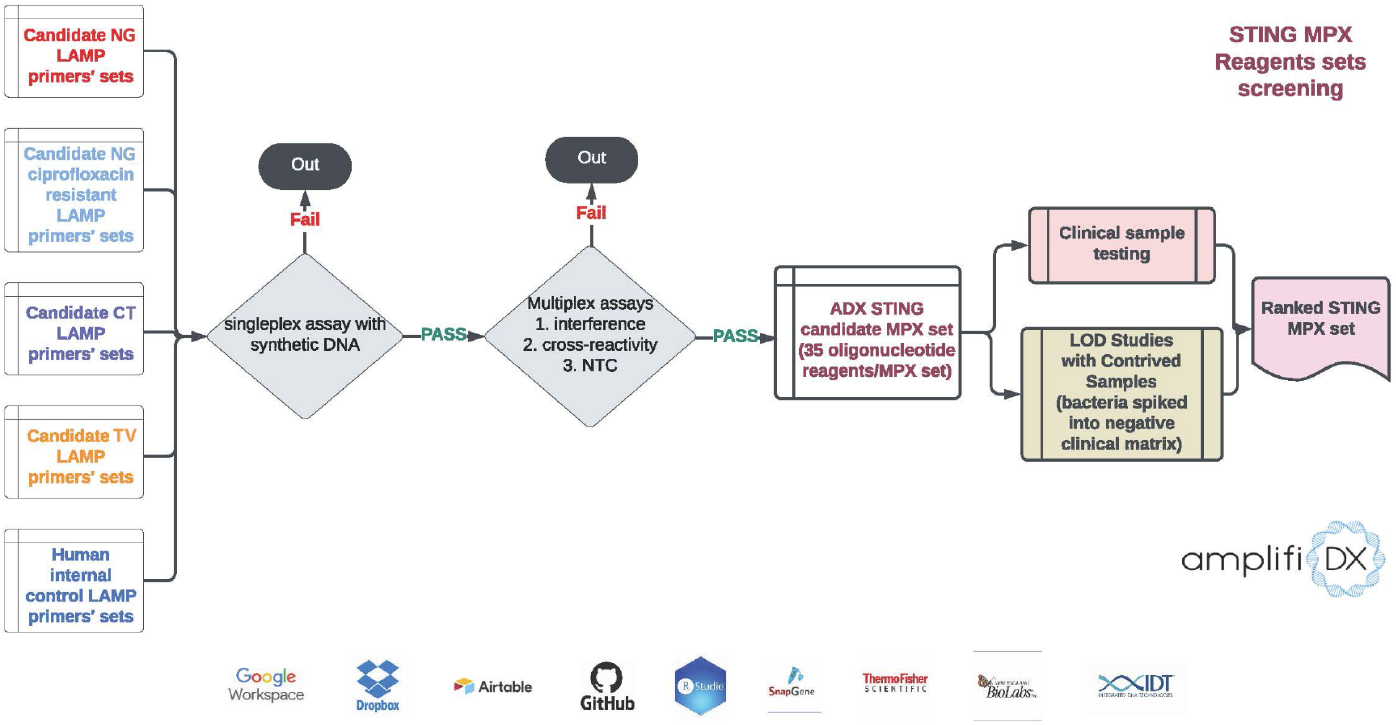
Workflow for empirical screening of LAMP primer sets. Primer sets that passed *in silico* inclusivity and specificity filters (Figure 1) were subjected to a multi-stage empirical screening process. First, primers were tested in singleplex reactions using synthetic gene fragments to confirm amplification efficiency and acceptable time-to-detection (TTD ≤15 min). Sets with consistent amplification were then transitioned into multiplex reactions using target-specific fluorescent probes. Multiplexed assays were evaluated for amplification efficiency, primer–primer interference, and background signal. Final primer sets with robust, reproducible performance were incorporated into the STI NG Plus assay chemistry.

Primer sets demonstrating efficient single-target amplification were advanced to multiplex screening using target-specific fluorescent probes. Multiplex formulations containing up to 35 oligonucleotides were assessed for amplification efficiency, absence of primer-primer interference, and reproducibility. Sets with robust performance and minimal background were incorporated into the final STI NG Plus chemistry used in this study.

### Organism Stock Quantification

Strains used in this study were obtained from ATCC or BEI and included: *Chlamydia trachomatis* (ATCC VR-348B) (CT), *Neisseria gonorrhoeae* (ATCC 49226) (NG WT), *Neisseria gonorrhoeae gyrA* S91F (BEI 1130991) (NG MUT), and *Trichomonas vaginalis* (ATCC NYCA04) (TV). Stocks of CT were diluted 500-fold in TE buffer to TCID50 ≥ 10. Stocks of NG WT, NG MT and TV were diluted 1:10 with the lysis buffer and used for digital PCR quantification. Dilutions were aliquoted and stored at -80°C. Aliquoted samples were then diluted in AmplifiDx lysis buffer at 1:1 ratio to lyse the cells and denature the double-stranded bacterial genomic DNA. Lysate was used directly for digital PCR quantification.

Digital PCR (dPCR) was performed using the Qiagen QIAcuity Four dPCR system (Qiagen cat no. 911042) according to the manufacturer’s instructions. For quantification of DNA, the QIAcuity EG PCR Kit (Qiagen cat no. 250111) was used with 1X master mix, 400 nM each forward and reverse primers, and 20% (v/v) final concentration of template. Each sample was run in duplicate using QIAcuity 8.5k 24-well Nanoplates (Qiagen cat no. 250011). 14 uL of dPCR mix was loaded into each well. Thermocycling was performed with an initial denaturing at 95 ^°^C for 2 min, followed by 40 cycles of 95 ^°^C for 15 sec, 60 ^°^C for 15 sec, and 72 ^°^C for 15 sec, with a final cool down at 40 ^°^C for 5 min. Plate images were obtained in the green channel with a 200 ms exposure and a gain value of 6.

### Sample Preparation and LOD Determination

Quantified stocks samples were prepared by diluting stocks into either clean STI NG Plus lysis buffer (1x TE, IDT cat no. 11-05-01-13) or clinical matrix to generate contrived samples for assay Limit of Detection (LOD) testing (**Figure 3**).

**Figure 3.**
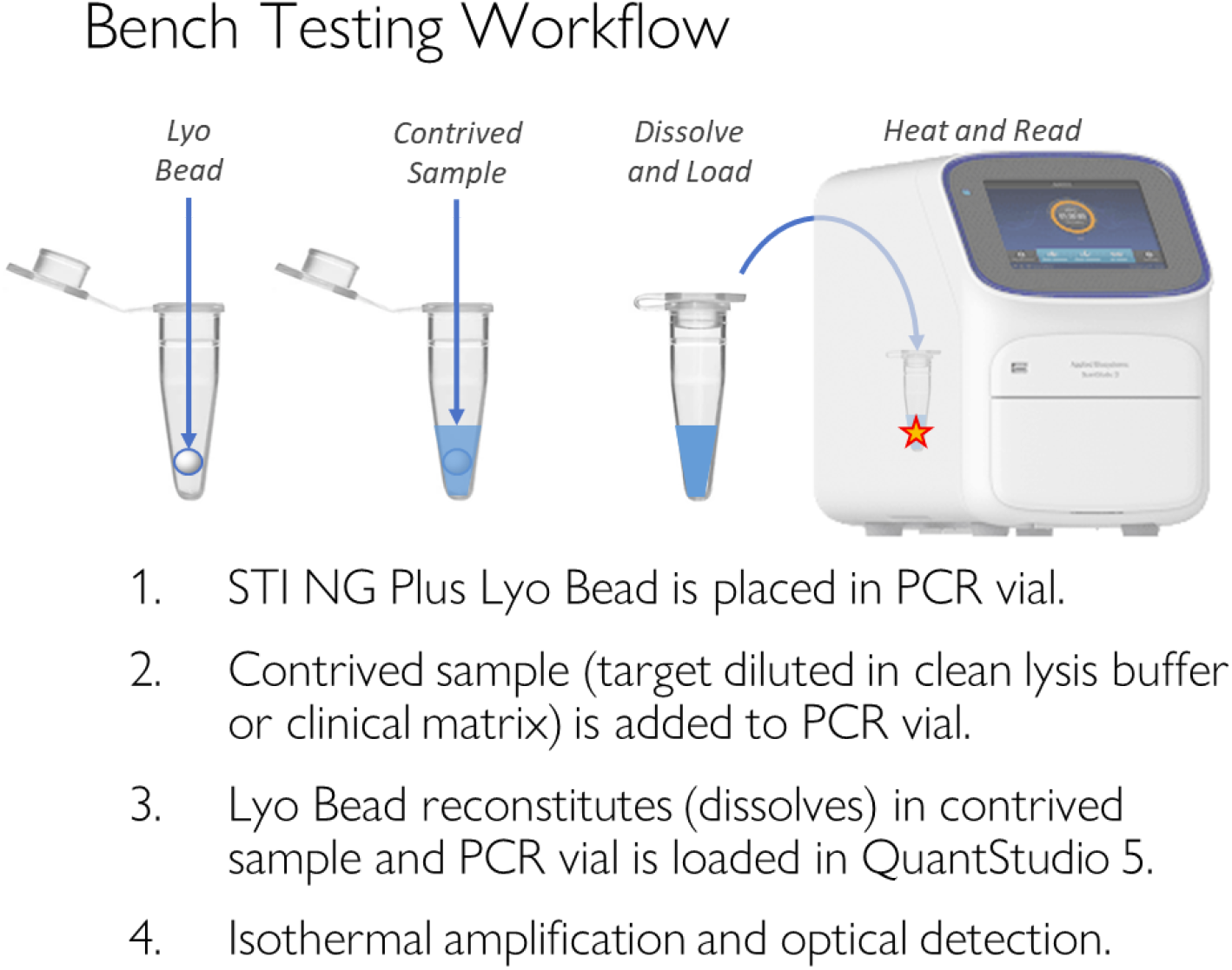
Schematic overview of the STI NG Plus testing workflow for bench testing, lyophilized reaction beads (Lyo Beads) were combined with contrived samples (spiked in clean lysis buffer or clinical matrix) in PCR tubes and loaded onto the QuantStudio 5 instrument for isothermal amplification and optical detection.

The LOD of the STI NG Plus assay was first evaluated using a clean buffer. CT and NG WT samples were initially prepared at concentrations corresponding to a FDA-cleared test LOD level: 755 GE/mL for CT and 245 GE/mL for NG. At each dilution level 10 replicates of each target were tested; if 10/10 were detected, targets were diluted 3-fold and testing was repeated until there was <95% detection. For TV and NG MUT microorganism testing, dilutions were initiated at a series of 10-fold, subsequently at 3-fold down from the last most diluted level of detection. At each dilution level, 3 replicates of each target were tested; if 3/3 were detected, targets were diluted 3-fold more and testing was repeated until there was <100% detection. Then, 10 to 12 replicates were tested at the previous highest dilution level with >95% detection.

LOD of the STI NG Plus assay was then evaluated with clinical matrix samples spiked with diluted microorganism stocks (**Figure 3**). Clinical matrix samples consisted of dry vaginal swabs sourced from University of Alabama at Birmingham (IRB Certificate of Action for Protocol #20232944). Vaginal swabs were from healthy donors, and had tested negative for all microorganisms included in the study via an existing cleared test. To prepare a clinical matrix, a swab was eluted into a 1x PBS buffer. Microorganism stocks were diluted with the swab eluate at the final targeted LOD level that was previously determined for the target microorganism in a clean buffer. The workflow was designed to be similar to the workflow that will be implemented in the POC device under development (**Figure 6**).

The negative control matrices from clinically negative swabs were used to confirm specificity. Samples were placed in PCR tubes preloaded with STI NG Plus amplification reagents. Samples underwent isothermal amplification and optical detection using a Thermo Fischer QuantStudio 5 Real-Time PCR instrument. Amplification and detection were performed at 63°C for 20 or 30 min and fluorescence was collected at 1 min intervals.

## RESULTS

*In silico* analysis confirmed that the STI NG Plus primer sets achieved high inclusivity across all known genome variants of each target organism. Specifically, all *C. trachomatis* primers aligned perfectly to all 321 GenBank whole genome records. For *N. gonorrhoeae*, 5 out of 351 genomes exhibited minor nucleotide variability in a subset of primers, but no mismatches were observed in critical regions required for amplification. For *T. vaginalis*, minor variability was observed in a single primer (LB) across 11 of 287 genomes, with all variation found in draft assemblies where sequencing artifacts could not be ruled out. These results indicate that the selected primers maintain comprehensive coverage of known pathogen diversity.

Cross-reactivity screening against 48 organisms commonly found in the urogenital tract, including *Gardnerella vaginalis, Mycoplasma genitalium, Candida albicans, Escherichia coli*, and *Lactobacillus* spp., demonstrated high assay specificity. No primer sets met alignment criteria predictive of off-target amplification. In addition, none of the primers exhibited significant alignment to the human genome (GRCh38), confirming absence of host interference. These findings provide confidence in the assay’s specificity and support the observed clean amplification profiles in both buffer and clinical matrix samples.

The STI NG Plus multiplex LAMP assay successfully amplified and detected DNA from all four target organisms—*Chlamydia trachomatis* (CT), *Neisseria gonorrhoeae* wild type (NG WT), *Neisseria gonorrhoeae* gyrA mutant (NG MUT), and *Trichomonas vaginalis* (TV)—in both clean lysis buffer and negative clinical matrix. Amplification curves generated using the QuantStudio 5 system demonstrated robust signal strength and consistent amplification kinetics across multiple replicates, supporting reliable detection at low template concentrations.

In clean lysis buffer, the assay demonstrated high analytical sensitivity, with limits of detection (LODs) substantially lower than those of the existing FDA-cleared test. Specifically, LODs for CT and NG WT were approximately 31-fold and 7-fold lower, respectively, compared to the FDA-cleared test (755 GE/mL for CT and 245 GE/mL for NG) (**Table 2**). Amplification was observed in all replicates for all pathogenic samples at low concentrations, confirming the efficiency and sensitivity of the LAMP amplification chemistry (**Figure 4**).

**Table 2.**
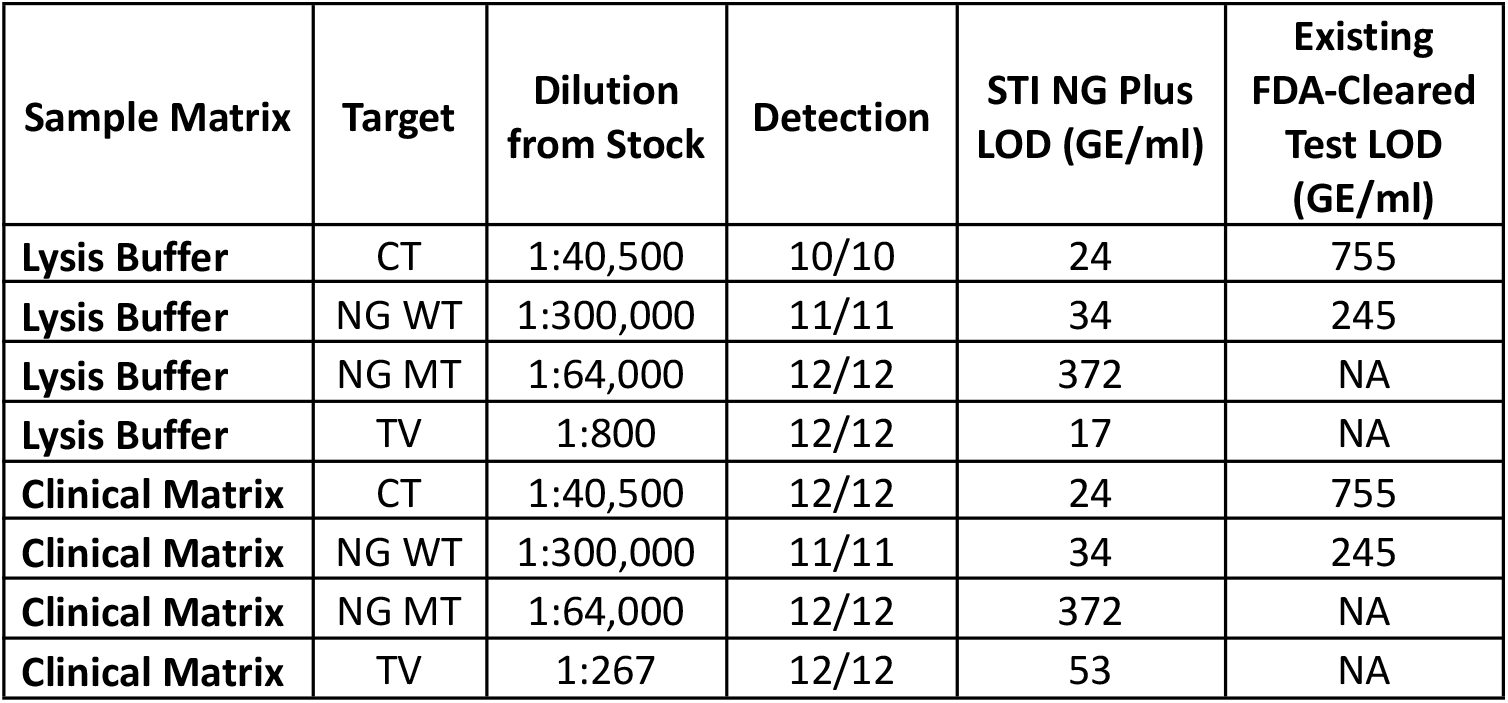
Analytical sensitivity results for the STI NG Plus Assay run with contrived samples in both lysis buffer and clinical matrix. Limits of detection (LOD) in genome equivalents per milliliter (GE/mL) for *Chlamydia trachomatis* (CT), *Neisseria gonorrhoeae* wild type (NG WT), *Neisseria gonorrhoeae* gyrA S91F (NG MUT), and *Trichomonas vaginalis* (TV) using the STI NG Plus assay. Results are shown for both clean lysis buffer and clinical matrix. The assay demonstrates lower LODs for CT and NG WT than an FDA-cleared test. Detection was confirmed in ≥95% of replicates at the listed concentrations.

**Figure 4.**
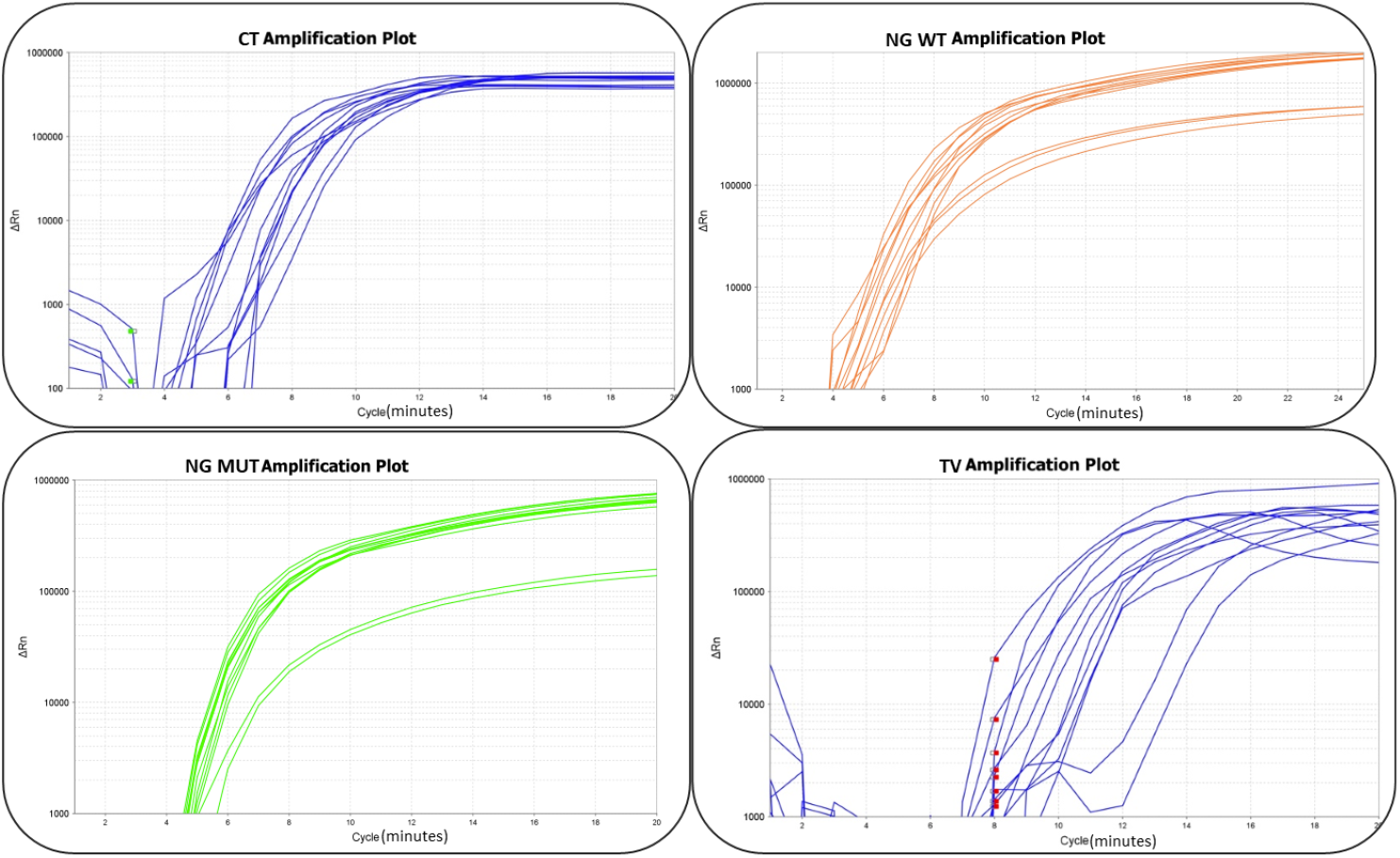
Amplification curves in clean lysis buffer matrix. Representative LAMP amplification curves for each of the four target organisms—*Chlamydia trachomatis* (CT), *Neisseria gonorrhoeae* wild type (NG WT), *Neisseria gonorrhoeae* gyrA mutant (NG MUT), and *Trichomonas vaginalis* (TV)—using quantified stocks spiked into clean lysis buffer. Reactions were run on the QuantStudio 5 platform. All targets amplified efficiently at low genome equivalent concentrations, with consistent signal profiles across replicates, demonstrating high analytical sensitivity and reproducibility under ideal reaction conditions.

Critically, the assay also demonstrated strong performance in clinical matrix samples derived from swab eluates from healthy patients. Pathogen DNA spiked into this matrix was consistently amplified and detected at low concentrations, validating the assay’s robustness in the presence of compounds found in patient samples (**Figure 5**). These findings support the feasibility of the assay chemistry using direct patient swab eluates without extraction or extensive sample processing.

**Figure 5.**
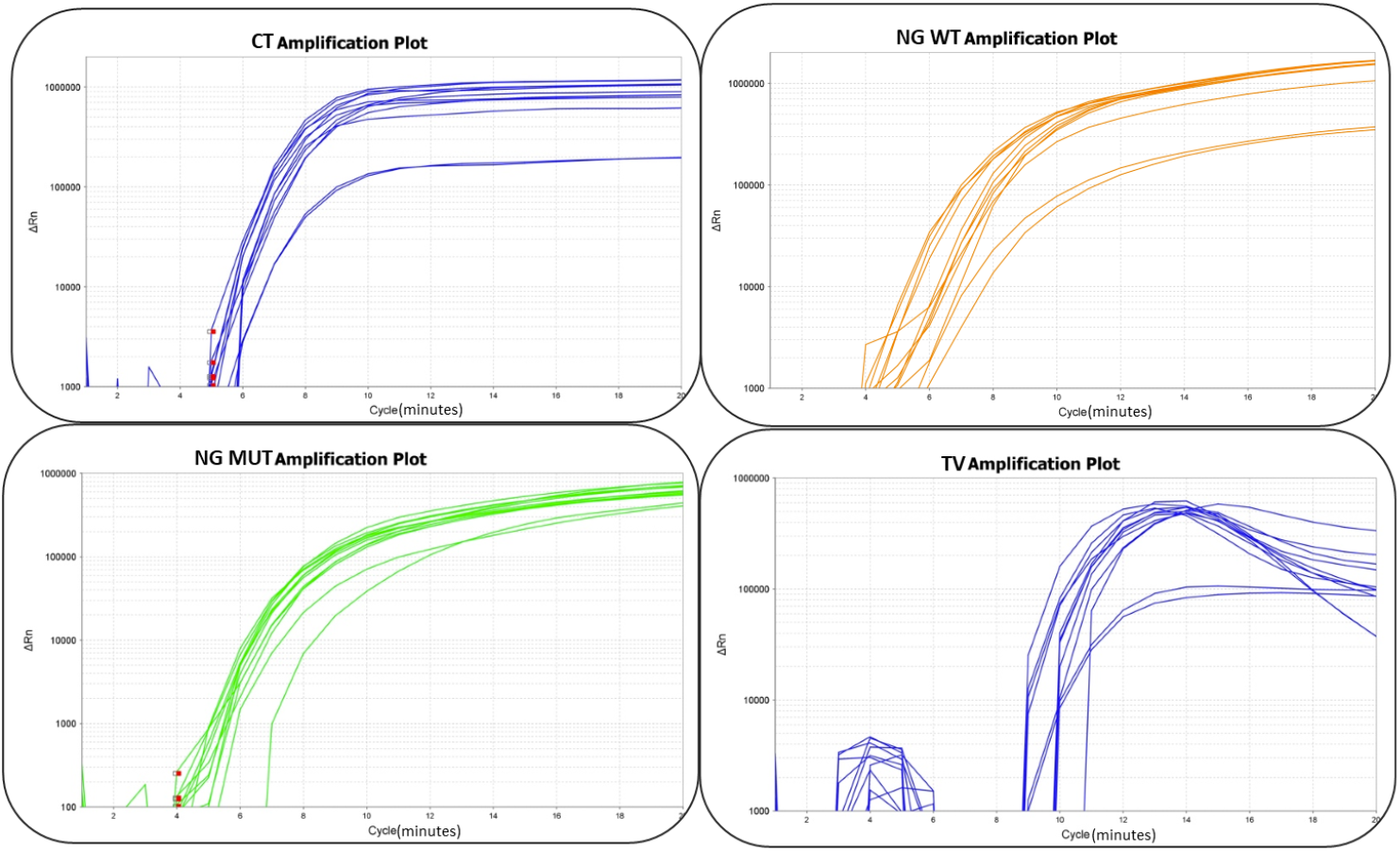
Amplification curves in clinical matrix. Representative LAMP amplification curves for each of the four target organisms—*Chlamydia trachomatis* (CT), *Neisseria gonorrhoeae* wild type (NG WT), *Neisseria gonorrhoeae* gyrA mutant (NG MUT), and *Trichomonas vaginalis* (TV)—using quantified stocks spiked into negative clinical matrix derived from swab eluates. Despite the presence of complex biological components, all target organisms were reliably detected at low concentrations, confirming the robustness of the assay chemistry with real-world clinical specimens.

Overall, the LAMP-based STI NG Plus assay chemistry delivered detection for CT, NG WT, NG MUT, and TV in both lysis buffer and in clinically relevant matrices. The LAMP-based STI NG Plus assay requires no sample preparation, and has a rapid time to amplification, highlighting its suitability for rapid, point-of-care applications.

## Discussion

We have developed a rapid molecular multiplexed assay utilizing LAMP technology, specifically tailored for point-of-care settings. This multiplexed test simultaneously detects CT, NG, TV, and fluoroquinolone resistance mutations in NG (S91F of *gyrA*), addressing critical gaps in current diagnostic offerings. Including TV alongside the more commonly tested CT and NG provides significant clinical benefits, as 1) TV infections are often asymptomatic yet associated with serious reproductive health consequences; 2) symptoms may mimic those of CT and NG; and 3) co-infection of TV with other STIs is high in vulnerable populations [23]. Early and simultaneous detection through multiplexing supports prompt, targeted treatment and clinical management, and reduces the risk of ongoing transmission and long-term complications [16,24,25].

The strong analytical performance of the STI NG Plus assay is supported by a rigorous *in silico* design and screening process. Primer sets were selected based on comprehensive inclusivity analysis across hundreds of whole-genome sequences for each target organism, ensuring detection of known genetic diversity, including variants of clinical and epidemiologic relevance. This is particularly important for pathogens like *N. gonorrhoeae* and *T. vaginalis*, where sequence variability can impact diagnostic accuracy. In addition, cross-reactivity screening against a diverse panel of urogenital flora and the human genome minimized the risk of false positives due to off-target amplification. This high level of target discrimination is especially critical in LAMP assays, which use multiple primers per target and are more susceptible to primer–primer interactions and non-specific signals if poorly designed. The combination of bioinformatic and empirical screening steps resulted in a robust multiplexed chemistry with excellent sensitivity, specificity, and reproducibility across clinical matrices.

The STI NG Plus assay demonstrated high analytical sensitivity, capable of reliably detecting low pathogen loads. Critically, the assay performed robustly in clinically derived sample matrices, confirming its suitability for realistic clinical scenarios. This robust performance underscores the assay’s potential to enhance clinical decision-making accuracy at the point of care [13-15].

Detection of NG gyrA-positive samples highlights another significant benefit of this multiplexed test. Identifying fluoroquinolone resistance markers enables healthcare providers to tailor antibiotic therapy effectively, thus promoting antibiotic stewardship. This targeted approach helps mitigate inappropriate antibiotic use and combat the rising threat of antimicrobial resistance in NG, preserving the efficacy of existing medications [10-12].

To further advance the assay’s clinical applicability, the next steps include transitioning to testing cartridges on the intended POC instrument, DX 100, currently under development (**Figure 6**), and evaluating assay performance with clinical patient samples against an accepted method. These steps will validate the assay’s practical effectiveness and readiness for clinical deployment.

**Figure 6.**
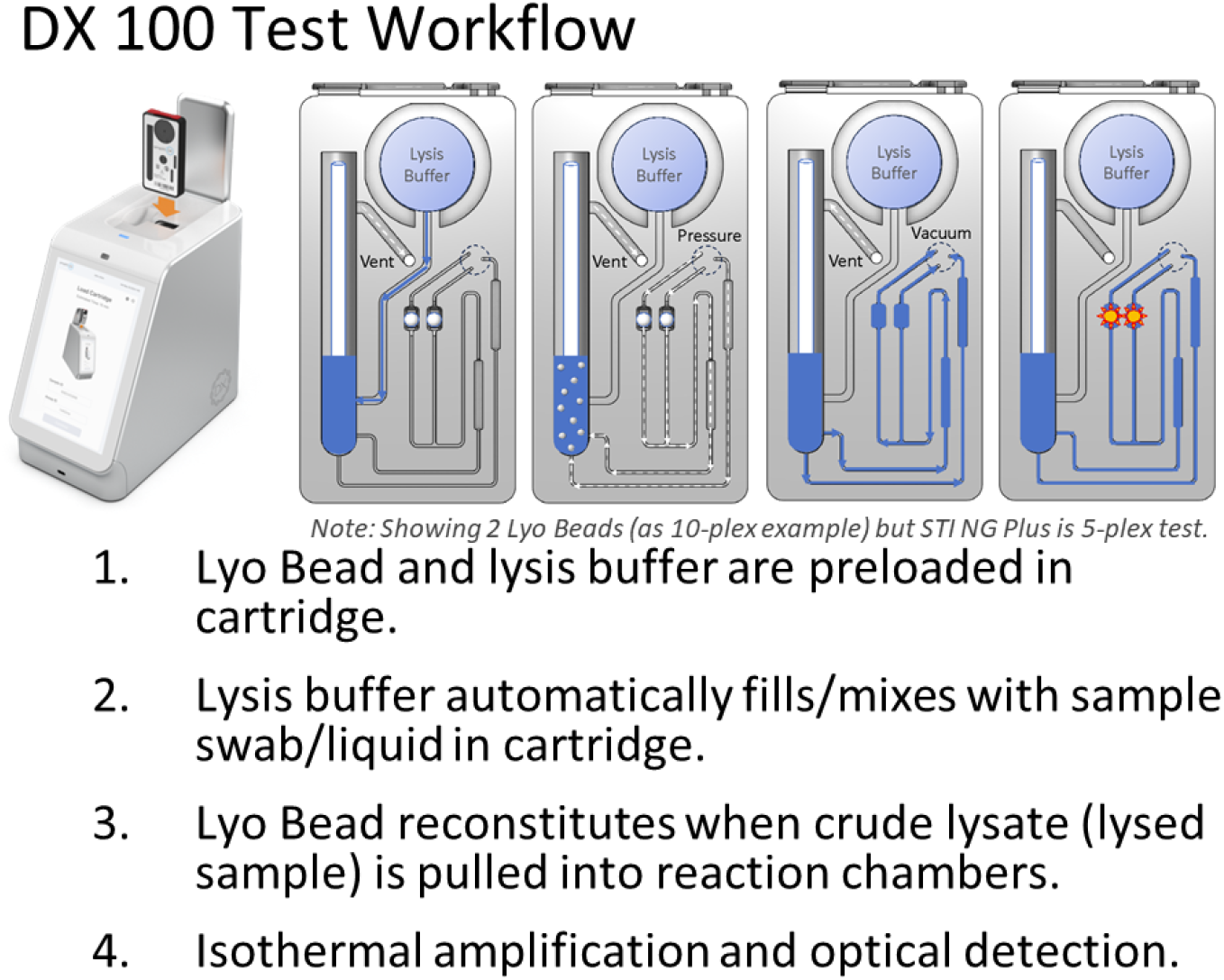
Proposed next phase of testing to be run on the DX 100 instrument, currently under development. The DX 100 platform will utilize a similar process as the bench testing (**Figure 3**) in a closed cartridge system. Briefly a dry swab or liquid sample is mixed with preloaded lysis buffer within the cartridge, reconstituting the lyophilized reagent bead (Lyo Bead) and initiating the amplification reaction within integrated chambers. Both bench and cartridge workflows use the same chemistry and amplification conditions.

Technological advancements, such as this multiplexed LAMP assay, have profound implications for patient care by enhancing accessibility, speed, and accuracy of STI diagnostics. Through improvements in point-of-care testing, healthcare systems can more effectively address the escalating rates of STIs, including CT, NG, and TV, thereby improving patient outcomes and public health [6,7,16].

## Ethics

Informed consent was obtained from all study participants. (University of Alabama Birmingham IRB Certificate of Action for Protocol #20232944)

## Data and materials availability

All data supporting the findings of this study are available within the manuscript or from the corresponding author upon reasonable request. Materials, including the STI NG Plus assay chemistry, are available for research use. For inquiries regarding access to materials or data, please contact Chris Meda at.

